# Direct Brain Recordings Suggest a Causal Subsequent-Memory Effect

**DOI:** 10.1101/2022.10.12.511606

**Authors:** Daniel Y. Rubinstein, Christoph T. Weidemann, Michael R. Sperling, Michael J. Kahana

## Abstract

Endogenous variation in brain state and stimulus-specific evoked activity can both contribute to successful encoding. Previous studies, however, have not clearly distinguished among these components. We address this question by analyzing intracranial EEG recorded from epilepsy patients as they studied and subsequently recalled lists of words. We first trained classifiers to predict recall of either single items or entire lists and found that both classifiers exhibited similar performance. We found that list-level classifier output—a biomarker of successful encoding—tracked item presentation and recall events, despite having no information about the trial structure. Across widespread brain regions, decreased low- and increased high-frequency activity (HFA) marked successful encoding of both items and lists. We found regional differences in the hippocampus and prefrontal cortex, where in the hippocampus HFA correlated more strongly with item recall, whereas in the prefrontal cortex HFA correlated more strongly with list performance. Despite subtle differences in item- and list-level features, the similarity in overall classification performance, spectral signatures of successful recall, and fluctuations of spectral activity across the encoding period argue for a shared endogenous process that causally impacts the brain’s ability to learn new information.

---

Fluctuations in neural processes during encoding contribute to the likelihood of recalling a given experience (Griffiths, Mazaheri, Debener, & Hanslmayr, 2016; deBettencourt, Norman, & Turk-Browne, 2018). Studies using functional neuroimaging and electrophysiological methods have demonstrated that neural activity measured during encoding of individual items reliably predicts their subsequent retrieval, an effect termed the subsequent memory effect (SME) (Wagner et al., 1998; Paller & Wagner, 2002; Sederberg, Kahana, Howard, Donner, & Madsen, 2003; Kim, 2011). However, key questions remain unanswered regarding the relationship of the SME to memory encoding states. First, most SME findings focus on individual items, where factors such as serial position, semantic characteristics, or idiosyncratic autobiographical associations may independently influence encoding success regardless of ongoing cognitive processes in the brain, leading to the possibility that the SME largely reflects these exogenous factors instead of memory-related internal states (Bainbridge, Hall, & Baker, 2019; Aka, Phan, & Kahana, 2021; Halpern, Tubridy, Davachi, & Gureckis, 2021). Second, the extent to which neural signals associated with SMEs reflect item-specific processing versus longer time-scale fluctuations in the brain’s ability to encode information, remains unknown. Thus, the question of whether the SME truly captures endogenously varying memory-related states, and how to more effectively measure such states, is unresolved.

Some studies have addressed the question of exogenous versus endogenous sources of encoding by controlling for temporal order effects known to predict recall (Serruya, Sederberg, & Kahana, 2014; Kahana, Aggarwal, & Phan, 2018; Aka et al., 2021; Weidemann & Kahana, 2021). Weidemann and Kahana (2021) show that even controlling for such variables, scalp EEG-recorded activity significantly predicts recall, suggesting that we can observe meaningful internal states with neural recordings. Consistent with the finding of endogenous factors underlying recall, Kahana et al. (2018) show that accounting for external predictors of recall such as alertness, temporal order, and a general measure of recallability, still leaves a large proportion of variability in performance unexplained—variability that we may be able to partially explain if neural activity can reveal internal states.

As Weidemann and Kahana (2021) demonstrate, one way to control for some of the confounds affecting memory, such as serial position, is to average neural activity and performance over multiple items. This multi-item analysis may also reveal different, longer time-scale aspects of the brain’s ability to encode information, as has been shown using fMRI in a study of state-related SME (Donaldson, Petersen, Ollinger, & Buckner, 2001; Otten, Henson, & Rugg, 2002). Findings of pre-stimulus SMEs in the hippocampus also hint at the importance of considering longer time periods in predicting memory performance (Park & Rugg, 2010; Urgolites, Wixted, Goldinger, Papesh, & Treiman, 2020). Since the brain cements episodic memories over extended periods of time, examining slightly longer-term SMEs may reveal unique neural mechanisms and systems, that contribute to encoding and integration of new memories into existing schema (Preston & Eichenbaum, 2013). Thus, investigating memory encoding across multiple studied items could give more complete insight into the physiology underlying encoding states, leading to a more accurate assessment of such states.

To this end, we constructed multivariate classifiers based on intracranial EEG recordings to predict recall over multiple items, and compared the performance of these classifiers to those predicting single item-level recall. We then investigated the temporal dynamics of classifier output to determine if multi-item level classifiers could reveal internal encoding states, acting as biomarkers of successful encoding. Finally we examined the important neural features of multi-item classifiers and single item-level classifiers, to better understand the neural underpinnings of good encoding states.

## Methods

From a pool of 259 total participants who performed either random or categorized free recall, we first selected 66 patients who performed both versions, to increase the amount of data for each subject and allow our results to generalize across task manipulations, although we did not account for task version in any analyses here. We then selected patients who recalled an average of at least one item per list, contributed data from at least 10 lists per session, and from at least 24 lists in total (i.e., across all sessions). This resulted in a sample of 62 patients. We trained two types of classifiers to predict either individual item recall or list-level recall performance, based on neural activity during word presentation or average activity over entire lists, respectively.

### Participants

All patients, who had medication-resistant epilepsy, provided informed consent to be enrolled in the Defense Advanced Research Projects Agency (DARPA) Restoring Active Memory (RAM) project. Patients underwent neurosurgical implantation of electrodes to identify and monitor seizure activity. During this time they also performed a variety of cognitive tasks. Data were collected across the following eight participating institutions: Columbia University Hospital (New York, NY), Dartmouth-Hitchcock Medical Center (Lebanon, NH), Emory University Hospital (Atlanta, GA), Hospital of the University of Pennsylvania (Philadelphia, PA), Mayo Clinic (Rochester, MN), National Institutes of Health (Bethesda, MD), Thomas Jefferson University Hospital (Philadelphia, PA), and University of Texas Southwestern Medical Center (Dallas, TX). Experimental protocols were approved by each Institutional Review Board.

### Free Recall Task

Patients performed two versions of a verbal delayed free recall task in which each session consisted of up to 25 lists with 12 words each. In one version, words were drawn randomly from a pool of 300 commonly used nouns (http://memory.psych.upenn.edu/WordPools). In the categorized version, words were drawn from a separate pool such that each list consisted of four words from three semantic categories (data previously published in Weidemann et al. (2019)). In each list, words were displayed for 1.6 s each, with randomly jittered inter-stimulus intervals of 0.75–1 s. Each word list thus lasted ~30 s. Following list presentation, patients performed a 20 s arithmetic distractor task of simple addition problems. Finally, patients had 30 s to recall as many words as possible. Patients completed as many sessions as was comfortable. The number of total sessions completed ranged from two to 13; 40 patients completed between two and four sessions, and 22 completed five or more sessions.

### iEEG Recording and Localization

We recorded neural activity using depth and surface electrode contacts. We constructed virtual bipolar contacts by subtracting the signal between adjacent monopolar contacts, and localized them to the midpoints of the two monopolar contacts. Monopolar contacts of a given bipolar pair were located within 20 mm of each other, outside of any clinician-determined seizure onset zone or region showing epileptic spikes. We registered post- and pre-implantation imaging using Advanced Neuroimaging Tools (ANTs) (Avants, Epstein, Grossman, & Gee, 2008). We localized surface contacts based on MRI segmentation using FreeSurfer (Desikan et al., 2006), and clinical neurophysiologists localized subcortical sources.

### iEEG Spectral Preprocessing

We aggregated recording segments from 0.3 to 1.6 s post word onset, for each list. We used Morlet wavelets (# cycles = 5) implemented in MNE-Python (Gramfort et al., 2013) to calculate spectral power at eight logarithmically-spaced frequencies from 3 to 180 Hz (3, 5, 10, 17, 31, 56, 100, 180) with a 1.2 s buffer period on each side of each segment. For each session and frequency, we log-transformed and z-scored the power. For list-level analyses we averaged power over all word presentation segments for each list. For analyses in which we applied the classifier to longer, continuous epochs (all 30 s before, during, and after list presentation, or the 4 s surrounding each item), we averaged power over the whole list presentation period, including inter-stimulus epochs, and after calculating power we down-sampled data to 500 Hz and averaged power over 1 s epochs incremented by 0.1 s.

### List-level Predictions

To predict performance of a given list, we implemented a linear regression model using the ‘sklearn’ ‘Ridge’ package in Python, with an L2 regularization parameter α of 1 /(2 * 0.0007), based on previously published results (Weidemann et al., 2019; Weidemann & Kahana, 2021). We trained models on all but one list and predicted recall performance on the held-out, test list (with each list held out once), using features of encoding epoch power at all eight frequencies, in all contacts (eight features per contact). To normalize power, we z-scored the training set of list-level power values within session, using the mean and standard deviation of the test list’s session to normalize the test list. To normalize list performance, we logit-transformed list performance, *p*, adjusting for performance of 0% or 100% (*p* = 0 or *p* = 1) by using: 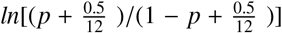 (Stevens, Valderas, Doran, Perera, & Kontopantelis, 2016). Otherwise we used the standard formula: *ln*[*p*/(1 – *p*)]. Next, we mean-subtracted list performances within each session, first omitting the held-out list’s performance to protect the training data from testing data. We then subtracted the mean performance of the corresponding session from the test list’s performance. Finally, we correlated predicted and observed list performance to quantify the overall performance of the model for a given subject. For parametric statistical tests, we used the Fisher transformation of the correlation values. We also performed permutation testing to obtain a distribution of 50 baseline correlation values by randomly shuffling list performances within each session and recalculating the correlation between predicted and observed list performance. Correlation values were deemed significant if greater than 95% of the baseline values.

### Item-level Classification

We performed item-level classification similarly to list-level prediction, except instead of using linear regression we used logistic regression, which is more appropriate for binary variables. We used L2-regularized logistic regression, implemented using the ‘sklearn’ ‘LogisticRegression’ package in Python, with a regularization parameter C of 0.0007, to be analogous with the L2 regularization performed in list-level predictions and to be consistent with prior work (Weidemann et al., 2019; Weidemann & Kahana, 2021). We set the “balanced” class weight parameter, which adjusts weights inversely proportional to the class frequencies, to account for the imbalance of recalled and unrecalled items. For each test list, we trained the model on all items from all other lists. To evaluate the classifier we compared observed recall to classifier-predicted probability of recall. We also calculated classifier-predicted list performance by summing the predicted probabilities of each individual item in the held-out test list. We constructed permutation-based baseline values similarly to in list-level classification.

### Recallability Correction

To correct for item and list recallability in item- and list-level prediction, we first measured item recallability as in Kahana et al. (2018), by calculating the probability of item recall within each patient and averaging these probabilities across patients. For each patient, we calculated this recallability measure for each item viewed based on all other patients. During recall classification, we first used this recallability measure to predict recall for all items excluding those in the held-out list, in a linear regression model. We applied this same model to items in the held-out list. We then used the residuals from these predictions as the new recall values, which we trained the linear regression models to predict, as above, based on neural activity measures. For the list-level analysis the individual item recallabilities were averaged over each list to generate list-level recallability measures.

### Shuffled List Control

To dissociate the confounding effects of item-level SME on list-level SME, we used the approach from Weidemann and Kahana (2021) and generated new lists for each subject according to the following procedure: within each session we aggregated the recalled and unrecalled items and randomly shuffled each of them, separately. For each list within each session, we picked recalled items from the shuffled set until the list contained the same number of recalls as in the true data. We then picked the remainder from the set of unrecalled items. We excluded those picked items from being chosen for subsequent lists; that is, items were picked without replacement. Finally, we repeated the list-level performance prediction procedure described above, 50 times, to generate 50 values of correlation between observed and predicted list-level performance. We used the mean of these 50 repetitions for each subject.

### Cross Classification

To assess the relative performance of the classifiers, we tested the item-level classifiers on list-level recall prediction, and vice versa. First, we trained the item-level classifier as above, and tested it on the neural features of the held-out list to generate the prediction of recall. We correlated these predictions with the observed list recall performance. To test the list-level classifier on item-level prediction, we trained the list-level classifier as above and tested it on the neural features of each item of the held-out list. We correlated these predictions of recall performance with observed item-level recall (a point-biserial correlation).

### Temporal Analysis of Encoding State

#### Temporal analysis of classifier output

We analyzed temporal fluctuations of classifier output by first training an item-level or list-level classifier as described above, except that for the training epochs of the list-level classifier we averaged power over the whole list instead of just word presentation epochs. For each test list, we applied the trained classifier to 1–second sliding windows, from 31 s before, to 61 s after onset of the first word of the list, incrementing by 100 ms. We also applied the classifier to windows from 2 s before to 2 s after individual words, to construct a peri-stimulus time course of classifier output. For both time courses, we averaged them for each subject to generate the subject-level mean.

#### Spectro-temporal analysis of classifier performance

We analyzed fluctuations in classifier performance over time and frequency using a similar temporal segmentation structure as above, but focusing only on the list presentation epoch (from 1 s before, to 31 s after onset of the first word), using 10 s time windows sliding by 1 s. For each time window we trained the list-level classifier on neural power from that time window alone and using only one of the eight frequencies, to predict performance on the entire list. We correlated these predictions of list performance with observed performances, over all lists, to quantify the classifier performance for that time-frequency cell. Similarly, we analyzed fluctuations in classifier performance over time alone, by training the list-level classifier on each 10 s time window using all eight frequencies.

### Correlation between Spectral Power and Performance

For each subject, we correlated both item- and list-level performance with power at each contact for each frequency.

We averaged correlations across all contacts for a given frequency to compare the item- and list-level correlations by frequency. To compare the item- and list-level correlations by region and frequency, we first aggregated contacts and averaged correlations over nine ROIs based on the grouping used in Weidemann et al. (2019): inferior frontal gyrus (IFG), middle frontal gyrus (MFG), superior frontal gyrus (SFG), temporal cortex (TC), hippocampus (HC), parahippocampal gyrus (PHG), inferior parietal cortex (IPC), superior parietal cortex (SPC), and occipital cortex (OC). We performed multiple comparisons correction for statistical tests using false discovery rate (FDR, *q* < 0.05) on permutation-based *p* values (Benjamini & Hochberg, 1995). We calculated *p* values by randomly shuffling region labels of electrode contacts 1000 times, at the subject-level, recalculating the group-level mean correlations for each permutation, and comparing these to the true mean correlation.

## Results

Our investigation addressed four main questions: 1) Can we reliably classify list-level recall performance and how does the performance of these classifiers compare with standard item-level prediction? 2) Do classifiers trained on items exhibit strong transfer to list-level recall, and vice versa? 3) Do fluctuations in predicted recall during encoding correspond with task-relevant events, thus enabling its use as a biomarker for encoding state? 4) Which aspects of neural activity—along the dimensions of time, frequency, and region—underlie recall prediction at the item and list level?

### Item- and List-level Classification of Memory Encoding

To evaluate list-level prediction of memory performance we first used a leave-one-list-out scheme to predict recall performance for each list, and then correlated the classifier-generated predictions of recall performance with observed performance (see *Methods* for details). To have a comparable measure for evaluation of the item-level classifier we correlated its predictions with the binary recall status of the corresponding items (i.e., a point-biserial correlation). At the individual subject level, list-level prediction was significant (permutation test, *p* < 0.05) in 40 of 62 patients. The distribution of correlations between predicted and observed list-level recall, across our 62 subjects, had a mean value of 0.25 (95% CI: [0.19, 0.31]) (*t*-test: *t*(61) = 8.8, *S E* = 0.029, *p* < 0.001). Item-level classification was significant in 60 of 62 patients, and the distribution of correlations for item-level recall had a slightly lower mean value of 0.22 (95% CI: [0.19, 0.24]) (*t*-test: *t*(61) = 18.2, *S E* = 0.012, *p* < 0.001). A statistical comparison of the item and list level correlations failed to detect any reliable differences (*M* = 0.027, paired *t*-test: *t*(61) = 1.44, *S E* = 0.027, *p* = 0.15). Figure 2 shows the comparable levels of performance of item- and list-level classifiers, and highlights the difference in variance.

**Figure 1.**
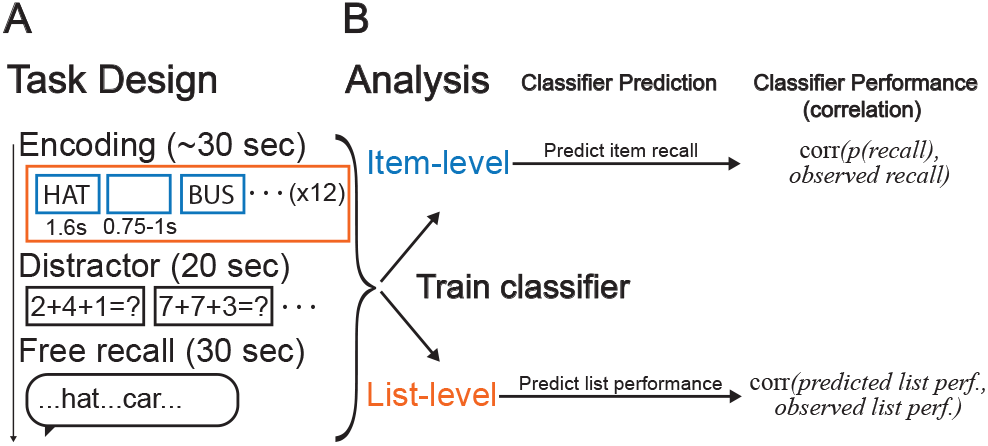
Task and Analytic Strategy. (**A**) Depiction of free recall task, consisting of up to 25 repeating blocks of encoding, distractor, and free recall epochs. (**B**) Depiction of analyses, where we generate item-level classifiers to predict probabilities of individual item recalls (*p*(*recall*)), and list-level classifiers to predict list-level performance. We evaluate classifier performance by correlating observed recall performance and predicted recall performance.

**Figure 2.**
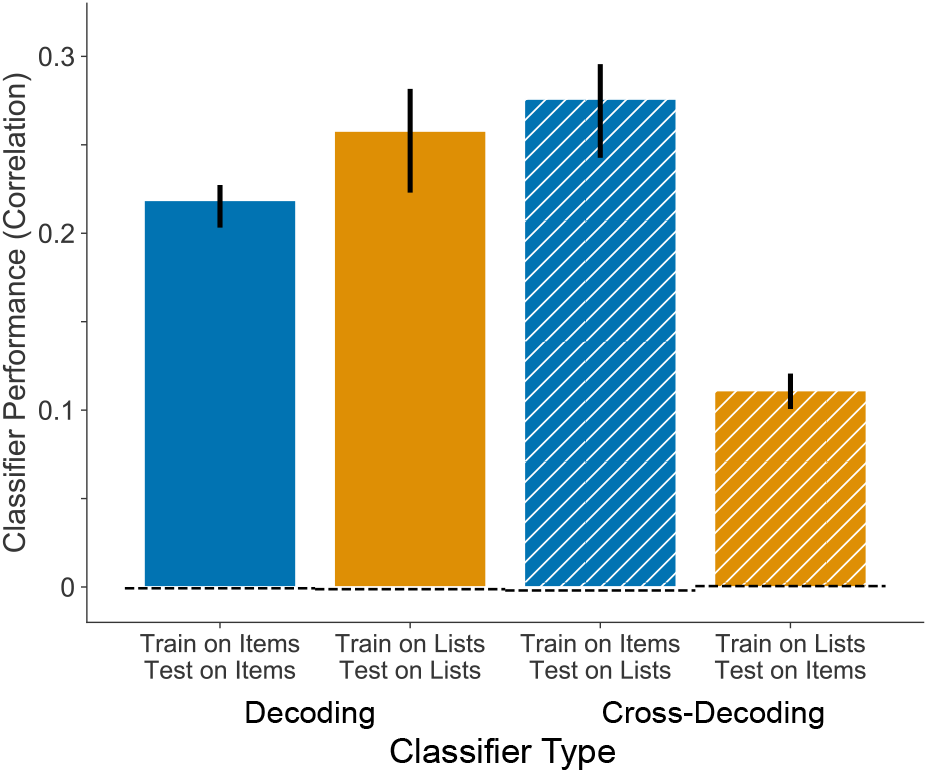
Classifier Performance. We quantified classifier performance as the correlation between predicted probability and observed recall (for item-level) or predicted and observed list performance (for list-level). We trained classifiers on either item-level recall (blue) or list-level performance (orange), and tested on either left-out items (outer bars) or left-out lists (inner bars). For cross-decoding, we trained the classifiers on item-level recall and tested them on left-out lists (hatched blue), or vice versa (hatched orange). We used permutation testing to derive the expected null correlation (dotted baselines), in which we randomly shuffled recall performances prior to classification for each subject (# permutations = 50). Error bars indicate ±1 SEM.

To verify that recall predictability was not solely due to item memorability effects, we repeated the above analysis but with first measuring and correcting for item recallability (Figure S3). After this correction, item-level classification was significant in 58 subjects, with a group mean of 0.17 (95% CI: [0.14, 0.19]) (*t*-test: *t*(61) = 12.8, *S E* = 0.013, *p* < 0.001). While still highly significant, these predictions were significantly decreased compared to no recallability correction (*M* = 0.049, paired *t*-test: *t*(61) = 11.3, *S E* = 4.5 × 10^-3^, *p* < 0.001). In contrast, list-level prediction of recall was not significantly affected by recallability correction (*M* = 3.7 × 10^-4^, paired *t*-test: *t*(61) = 0.60, *S E* = 3.2 × 10^-3^, *p* = 0.55). List-level classification after recallability correction was significant in 39 subjects, with a group mean of 0.25 (95% CI: [0.19, 0.31]) (*t*-test: *t*(61) = 8.7, *S E* = 0.030, *p* < 0.001).

To dissociate the confounding effects of item-level SMEs from the list-level, which is necessarily composed of items, we synthesized new lists while maintaining the true number of recalled items in each list. If the list-level performance predictions are significantly reduced, this would imply there is useful information at the list-level that is not found at the item-level. We found this to be the case, that shuffling lists significantly reduced the correlation between observed and predicted list-level performance in 56 of 62 subjects, and with a group mean difference of −0.15 (95% CI: [−0.20, −0.10]) (paired *t*-test: *t*(61) = −6.2, *S E* = 0.025, *p* < 0.001).

Item-level classifiers exhibited significant transfer to list-level predictability, and vice versa (i.e., cross-decoding) (Figure 2, hatched bars). Classifiers trained on item recall and tested on list performance exhibited a mean correlation of 0.27 (95% CI: [0.22, 0.32]) (*t*-test: *t*(61) = 10.4, *S E* = 0.026, *p* < 0.001). Prediction of item recall by classifiers trained on lists showed the lowest mean correlation value of 0.11 (95% CI: [0.09, 0.13]) but was still significantly positive (*t*-test: *t*(61) = 11.1, *S E* = 0.010, *p* < 0.001). Figure 2 shows that when predicting item recall, item-level classifiers performed better than list-level classifiers (paired *t*-test: *t*(61) = 11.0, *S E* = 0.010, *p* < 0.001). However, when predicting list performance, list-level classifiers did not perform better than item-level classifiers (paired *t*-test: *t*(61) = −1.02, *S E* = 0.018, *p* = 0.31). Overall, these results demonstrate that list performance is predictable to a comparable degree as single item recall, and given the cross-decoding success, the underlying neural features of item-level and list-level encoding states may largely overlap.

### Temporal Dynamics of Classifier Predictions

We next asked if classifier output could serve as a biomarker for encoding state. That is, would the temporal dynamics of classifier predictions of recall correspond to task-relevant events, and how would these dynamics differ between the two classifiers trained on item- and list-level recall performance? From the cross-decoding results showing generalizability between item-level and list-level classifiers, we hypothesized that the output of the two classifiers would display similar slow but not item-level temporal patterns, since the list-level classifier has no access to this scale of neural activity. However, Figure 3 shows that both the itemlevel and list-level classifiers exhibit similar item-level temporal fluctuations, peaking at 800 and 900 ms after word onset, respectively (Figure 3C&D). We also observed similarities in more slowly varying encoding state, such that both increased with onset of list presentation, distractor phase, recall period, and countdown period (Figure 3A&B). However, we also observed subtle differences at this temporal scale. Although solely descriptive and not a statistical analysis, itemlevel classifier predictions were highest during the first item and declined rapidly over the first half of the list, whereas list-level classifier output peaked at the second item of the list and was flatter over time.

**Figure 3.**
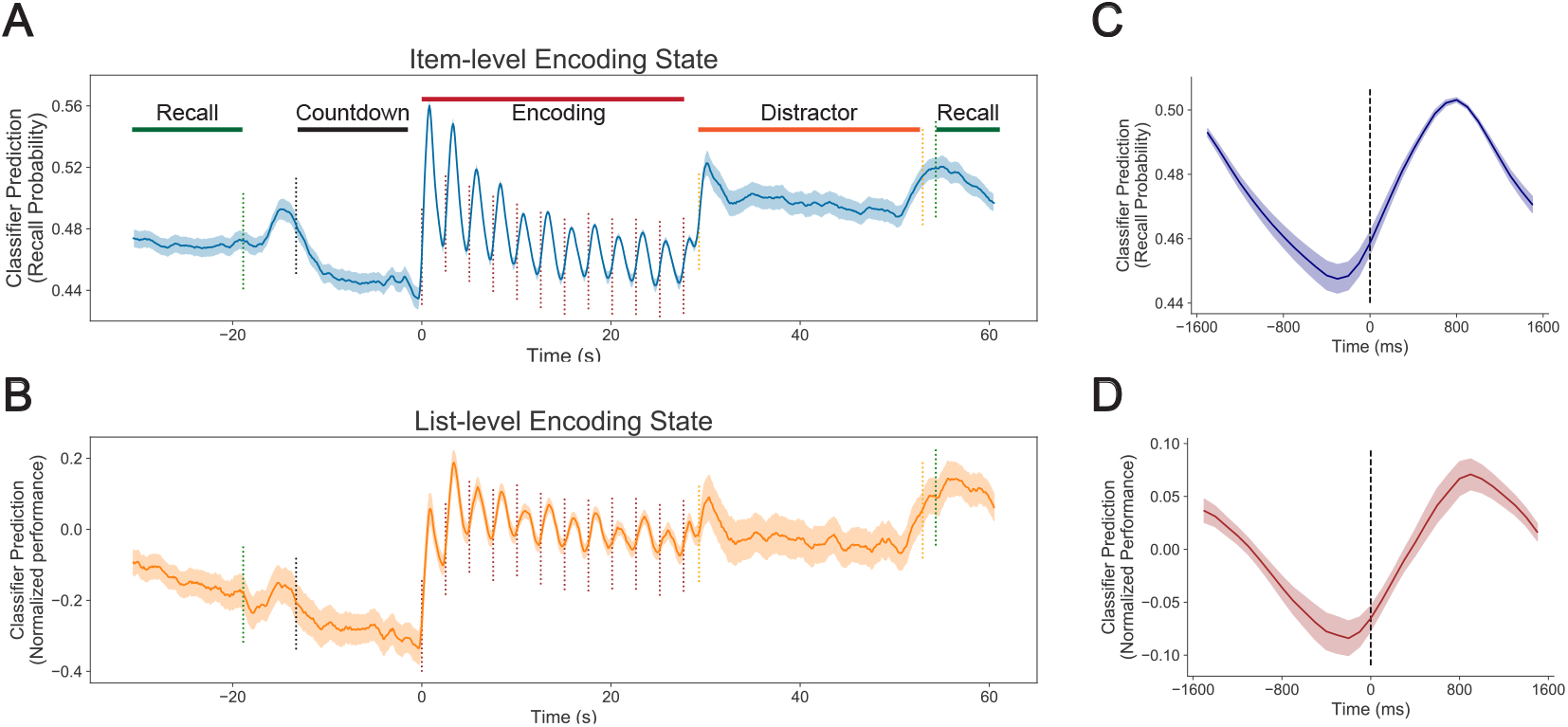
Classifier Predictions. (**A, B**) Time course of item-level (A) and list-level (B) classifier predictions over the course of encoding lists, showing correspondence with task-relevant events. We normalized list-level classifier prediction by subtracting the test list session’s mean performance. Dotted lines indicate average time of task events including start of countdown before list (black), word presentation times (brown), start/end of math distractor task (orange), and start/end of recall period (green). (**C, D**) Peri-stimulus time course of classifier predictions for item-level (C) and list-level (D) classifier, time-locked to word presentation time, highlighting peak classifier predictions at 800 ms (item-level) and 900 ms (list-level). All shaded error regions indicate ±1 SEM.

These results show temporal fluctuations correspond to task-relevant events and phases, and suggest that classifier output may be used as a biomarker of internal encoding state. Despite broad similarities between item- and list-level classifier outputs, especially at the single item level, minor differences suggest they reveal unique aspects of the encoding state.

### Physiological Substrates of Classification

With evidence that the classifiers reflect internal encoding states, we next examined the neural activity supporting classification. First, were there certain times during list presentation that were more useful in predicting whole-list performance? Given that recall of early-list items better predicts overall list performance (Figure S4), we hypothesized that early-list time windows of neural activity would also allow for better list-level recall predictions. However, list-level classifiers performed significantly better using the last time window compared to the first window (paired *t*-test: *t*(61) = 2.00, *S E* = 0.031 *p* = 0.0498); Figure 4A shows that this increase was relatively constant over the course of the list. The pattern of increasing predictability over time was generally common across frequencies but was particularly strong in alpha and high gamma frequencies (Figure 4B). Thus, neural activity at the end of the list may contribute more to list-level classification compared to activity at the beginning.

**Figure 4.**
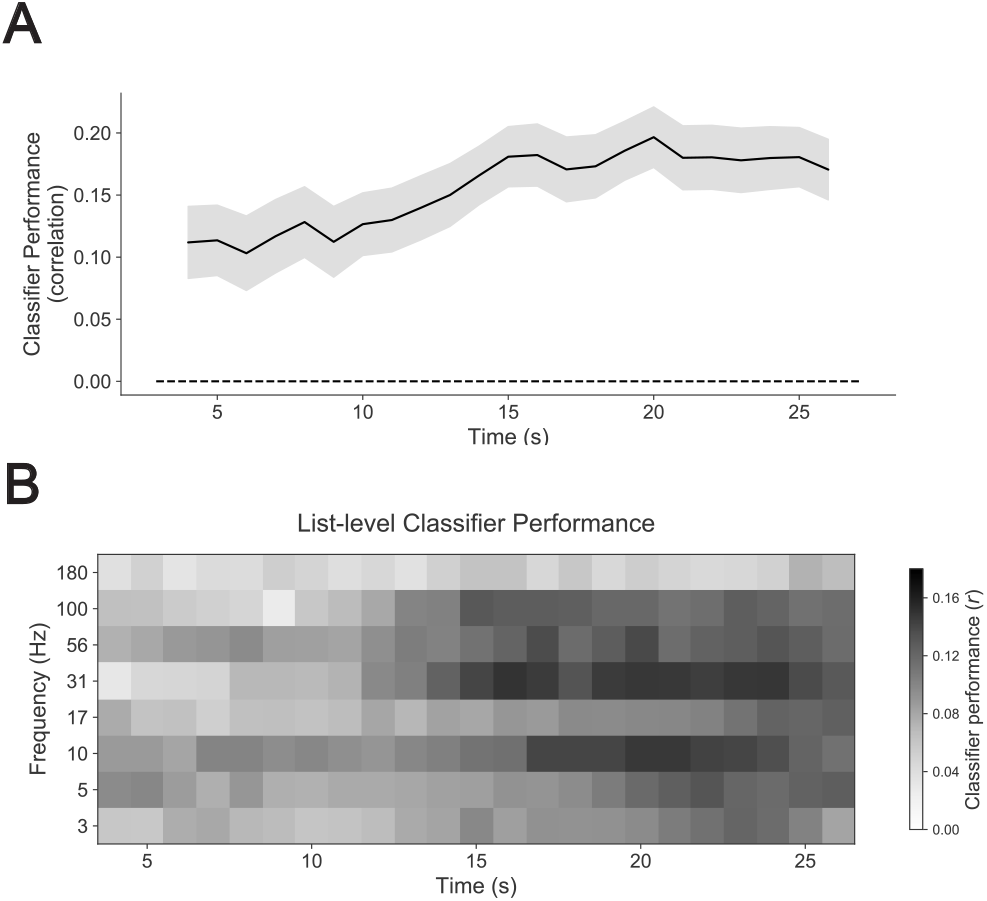
Spectro-Temporal Analysis of List Performance Prediction. (**A**) Representation of most useful times and frequencies for prediction of list performance. For each time point, we calculated spectral power in the 10 s time window centered on the time shown, relative to list onset. We then predicted list performance using power at this time point, with leave-one-list-out crossvalidation. List-level classifiers perform significantly better when using the end of the list compared to the beginning (*p* < 0.05). (**B**) Same as (A), except instead of using all frequencies together, we used one frequency at a time to predict list performance.

In addition to investigating the temporal aspects of the neural basis of multi-item encoding states, we investigated their spectral and regional aspects. Item- and list-level correlation patterns were similar, with the greatest negative correlation in the theta/alpha range, and the greatest positive correlation in the high gamma range (Figure 5A). The greatest item-level correlations between performance and power were at 5 Hz (*M* = −0.026, 95% CI: [−0.034, −0.018]) and 100 Hz (*M* = 0.016, 95% CI: [0.011, 0.020]). The greatest list-level correlations were at 10 Hz (*M* = −0.054, 95% CI: [−0.067, −0.027]) and 56 Hz (*M* = 0.021, 95% CI: [0.005, 0.036]). Correlations were significant with Bonferroni correction for multiple comparisons (uncorrected p’s < 0.002), except for item-level correlations at 31 Hz, and list-level correlations at 31 Hz and above.

**Figure 5.**
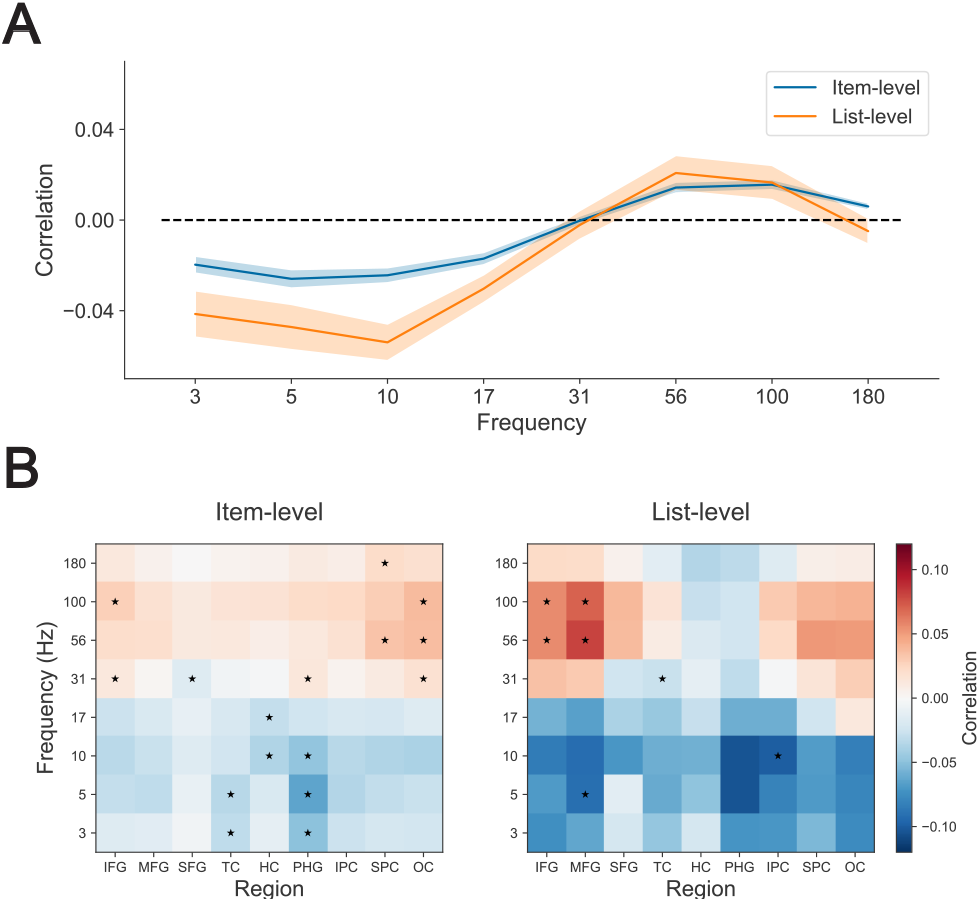
Univariate Correlations between Power and Performance. (**A**) We demonstrate a spectral tilt effect for itemlevel (blue) and list-level (orange) correlations between brain-wide power and recall performance at various frequencies. Shaded area around lines indicates ±1 SEM. (**B**) Same as (A) except here we report correlations by region of interest (IFG: inferior frontal gyrus; MFG: middle frontal gyrus; SFG: superior frontal gyrus; TC: temporal cortex; HC: hippocampus; PHG: parahippocampal gyrus; IPC: inferior parietal cortex; SPC: superior parietal cortex; OC: occipital cortex). Asterisks in cells denote statistical significance as determined by FDR correction at *q* < 0.05 after regional permutation analysis.

Finally, we computed correlations between performance and frequency-specific power for specific regions of interest. Item- and list-level correlation patterns by region were similar, with low-frequency negative correlations in parahippocampal regions and higher positive correlations in frontal and occipital regions. However, in contrast to item-level correlations, list-level correlations were relatively higher in frontal regions and exhibited less spectral tilt in temporal regions, especially in the hippocampus (Figure 5B). We specifically tested the difference between item- and list-level correlations in the hippocampus and DLPFC (middle frontal gyrus) in a subset of patients who had electrode contacts in both regions (*n* = 36) and found that univariate correlations between hippocampal HFA and recall performance were higher at the item-level compared to the list-level (paired *t*-test: *t*(35) = 2.28, *S E* = 0.014, *p* = 0.029). Conversely, correlations between prefrontal HFA and recall were higher at the list-level (paired *t*-test: *t*(35) = 2.52, *S E* = 0.011, *p* = 0.016) (Figure 6). Correlations between hippocampal HFA and recall performance were significantly positive at the item-level with mean of 0.010 (95% CI: [0.0008, 0.019]) (*t*-test: *t*(35) = 2.21, *S E* = 0.0044, *p* = 0.033), and while numerically negative at the list-level with mean of −0.021 (95% CI: [−0.052, 0.0097]), they were not significantly different from 0 (*t*-test: *t*(35) = −1.39, *S E* = 0.015, *p* = 0.17). Correlations between prefrontal HFA and recall were significantly positive at both the item-level (*t*-test: *t*(35) = 2.29, *S E* = 0.0047, *p* = 0.028) and the list-level (*t*-test: *t*(35) = 3.05, *S E* = 0.013, *p* = 0.0044) with means of 0.011 (95% CI: [0.0012, 0.020]) and 0.040 (95% CI: [0.013, 0.066]), respectively. Thus, while the broad physiological phenomenon of spectral tilt was apparent in both item-level and list-level SMEs, some specific patterns of regional contributions differed between the two temporal scales.

**Figure 6.**
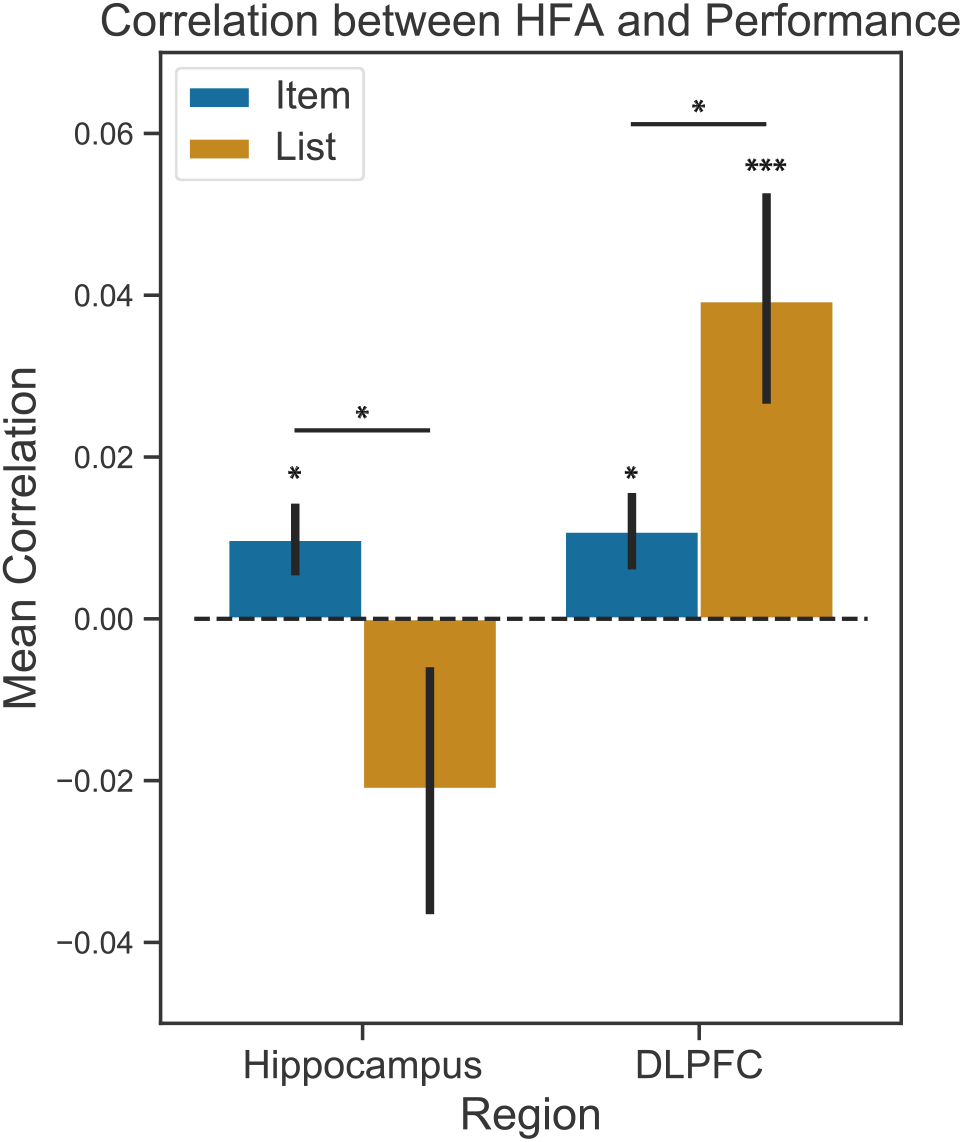
DLPFC and Hippocampal Contribution to Item-level vs. List-level Encoding. We averaged correlations between power and recall performance over all contacts within a given region (hippocampus: left; DLPFC or middle frontal gyrus: right) and all high gamma frequencies (56 Hz, 100 Hz, 180 Hz) for both item-level (blue) and list-level (orange) correlations, for each subject with contacts in both regions (*n* = 36). Hippocampal list-level correlations were not significantly different from 0 (*p* = 0.17). Paired *t*-tests reveal significantly greater item-level correlation with performance in the hippocampus, and greater list-level correlation in the DLPFC. Error bars indicate ±1 SEM. * denotes *p* < 0.05, *** *p* < 0.005.

## Discussion

The question of whether SMEs reflect causal internal encoding state has been unresolved due to confounding factors of item-specific characteristics. We approached answering this question by examining a longer-term SME over the multiple items in a list, and examining the time course of predicted encoding, using multivariate classifiers based on intracranial recordings. We demonstrate that this approach not only controls for some item-specific aspects, but also establishes a biomarker of encoding that reveals temporal dynamics of endogenous state. We also provide a more complete understanding of the SME in general by highlighting the similar and complementary regional contributions to SMEs at different time scales.

Whether using a short window of time to predict encoding success of a single item, or a long time window to predict encoding of multiple items, we found similar performance of encoding prediction. Performance was qualitatively higher using lists compared to single items, but not significantly so. Previous work employing a nearly identical free recall task with scalp EEG also found that predictions of recall from list-based classifiers correlated slightly better with observed performance than item-based classifiers, although a direct comparison was not performed (Weidemann & Kahana, 2021). While that study sampled healthy volunteers instead of patients with epilepsy, SMEs in both populations follow similar patterns (Hill, King, Lega, & Rugg, 2020).

One factor that complicates our direct comparisons of item-level and list-level classification is the large disparity in data quantity: item-level classifiers have 12 times the observations as the list-level classifiers, which affects the variance of the classification performance. We can somewhat account for the data disparity by repeating item-level classification analyses with smaller subsets of serial positions. Dividing the item-level classification analyses into four subgroups of serial positions (1–3, 4–6, 7–9, and 10–12) we find that the average correlations between predicted and observed recall drops from 0.22 to 0.14 (Figure S1). There was no significant difference between the average correlation of any of the individual subgroups, which ranged from 0.13 to 0.15 (p’s > 0.10). Although those subgroup-based classifiers have multiple times more data than the list-level classifiers, list-level classification correlations are likely higher due to the higher signal-to-noise ratio resulting from averaging features over longer time periods. With more data, list-level classifiers might be yet more accurate —when recalculating list-level classification performance using variable numbers of sessions in a subset of patients with the greatest number of sessions recorded, we find that classifier performance steadily increases from two to five sessions (Figure S2).

The success of list-level classification of memory performance is consistent with, and related to, previous studies of pre-stimulus and state-related SMEs (Donaldson et al., 2001; Otten et al., 2002), especially in the hippocampus (Urgolites et al., 2020; Park & Rugg, 2010), although some suggest that this signal is only relevant for recognition memory and not free recall (Merkow, Burke, Stein, & Kahana, 2014). Nevertheless, both pre-stimulus and list-level classification support the notion that neural activity outside of item presentation can still reliably predict memory encoding. With respect to list-level classification, previous work has shown that the effect is not driven solely by the predictability of the constituent items, since rearranging the items into new lists abolishes the ability to predict list-level performance (Weidemann & Kahana, 2021). To dissociate the confounding effects of item- and list-level SMEs, we replicated this analysis from Weidemann and Kahana (2021) and also found that shuffling lists significantly reduced the correlation between observed and predicted list-level performance in 56 of 62 subjects, with a group mean difference of −0.15 (95% CI: [−0.20, −0.10]) (paired *t*-test: *t*(61) = −6.2, *S E* = 0.025, *p* < 0.001). Relatedly, we retrained the list-level predictions model using only the inter-stimulus intervals (700 ms preceding word presentations) instead of word presentation times, and recalculated the time series of encoding prediction over time. While the resulting time series is noisier than when trained on the presentation time windows, the dynamics are overall the same (Figure S6). This suggests that the temporal dynamics of list-level predictions in Figure 3B follow item presentation timing not simply due to being trained on item-presentation windows. We also found that item-level classifiers generalized to predict list-level performance and vice versa (Figure 2), further suggesting that neural predictors of encoding success vary slowly.

### Causal vs. Non-causal SME

One major question arising from the SME studies has been whether correlations between neural activity and recall are causal, or rather represent correlations with external factors (Halpern et al., 2021) such as item memorability (Bainbridge et al., 2019), serial position (Murdock, 1962), or semantic characteristics across items in a list (Aka et al., 2021). We argue here that external characteristics of the words cannot be the only factor underlying prediction of recall, for multiple reasons. First, even when correcting for recallability, we find significant predictive success of the classifiers (Figure S3). Item-level classification success was significantly reduced, suggesting some contribution of item-specific effects to the item-level SME, however average correlations between predicted and observed recall was still highly significant. Furthermore, controlling for recallability did not affect list-level classification, confirming our hypothesis that averaging over individual items would abolish some effects of item-specific characteristics.

A second confound that may theoretically underlie item-level classification is serial position, but this cannot account for the list-level classification because the training data contains no serial position information. Furthermore, despite the stronger association of *early* item recall with whole-list performance, compared to later item recall (Figure S4), time segments *later* in the list provide the list-level classifier with better predictive information than time segments early in the list. This implies that predictors of list-level performance are not simply making use of aggregate item-level signals.

The inference that the SME identifies neural signals that specifically impact successful encoding of long-term memories, has recently been called into question. Specifically, Halpern et al. (2021) argue that well-known covariates of successful memory, such as serial position effects or item difficulty or linguistic properties that make certain items more memorable than others, may actually underlie previous studies claiming to identify neural correlates of success memory encoding. They report an fMRI study that controls for these variables that finds no evidence for SME in regions where it had previously been found. Here, by establishing list-level correlates of successful memory that remove any effects of serial position, and by showing that these effects remain robust even after controlling for item-level differences in word memorability, we see evidence for a causal SME.

However, as our study is inherently correlational, we cannot draw strong causal inferences about the specific brain states that lead to better memory. For example, variables that we have not controlled may lead to the observed brain states that are correlated with subsequent memory. Similarly, the observed SMEs may lead to other states that more directly cause memory formation. Although absolute causality is therefore impossible to prove, we have eliminated item-specific characteristics as the entire explanation of the SME, and therefore conclude that the neural SME used by the classifier here is likely at least partly causal.

### Encoding State Dynamics

Our finding that the temporal dynamics of the classifier output clearly coincide with task phases, supports the notion that the classifier output reveals internal encoding state. While the oscillations of classifier prediction with word presentation may possibly be related to item-level characteristics, the rise in prediction at the beginning of the distractor period, and especially during the recall period where there is no external stimulus, strongly suggest association with internal states untied to exogenous semantic measures. Furthermore, classifier predictions during recall increase more in high-performing lists (Figure S5). This is not merely a continuation of higher predictions during the encoding phase, as predictions during the distractor phase are similar between low- and high-performing lists. The oscillations with word presentation were particularly notable for the list-level classifier. One expects the item-level classifier to exhibit such dynamics since the training window is limited to the item presentation window, however the list-level classifier is trained only on the average power over the whole list, including the inter-stimulus windows, and still displays the identical pattern of encoding state peaks at 800–900 ms post word onset. This time course is consistent with previous findings that HFA-based SME peaks at around 700 ms, depending on the region (Sederberg et al., 2003; Burke et al., 2014). The two classifiers differed, however, regarding serial position effect. The item-level encoding state exhibited a dramatic serial position effect, reminiscent of the shifts in power observed by (Serruya et al., 2014) that also predicted subsequent recall. However, the list-level classifier exhibited a more subtle serial position effect, where encoding state peaked at the second item instead of the first, and declined to a lesser extent.

The rapid decline of the item-level classifier prediction may reflect attention-related processes or neural resources that gradually fatigue over time, and are renewed after a short break (Serruya et al., 2014; Tulving & Rosenbaum, 2006). During high-performing lists as well as low-performing lists, classifier prediction of item-level recall increases at the start of the distractor period (Figure S5). In contrast, only in low-performing lists does the list-level predicted recall increase at the start of distractor periods, suggesting that during higher memory performance (perhaps related to greater contextual clustering) list-level predictions are more memory-specific. More concrete modeling studies will further quantify these descriptive interpretations. Overall though, the time courses of item-level and list-level encoding states both appear to reflect times of increased task engagement, with transient rises at task-relevant moments.

We also examined the list-level classifier performance over time, to test if certain times during list presentation were more useful in predicting whole-list performance, and thus understand how individuals encode over time. Figure 4 suggests that items are not simply encoded sequentially, with each item’s encoding ceasing with the end of its presentation window. Instead, encoding of early items may continue in time such that by the end of the list, all preceding items are experiencing some degree of simultaneous encoding. Although speculative, this notion is also consistent with the phenomenon of rehearsal (Corballis, 1969), and that by the end of the list there are more items to potentially rehearse than at the beginning. Therefore, while speculative, there may be more list-level relevant information at the end of the list than at the beginning.

A previous investigation of encoding state based on pupil size produced a remarkably similar time course to that shown in Figure 3A&B (Kucewicz et al., 2018). Pupil size increased more during words that were recalled than unrecalled, and also during the retrieval period, demonstrating that pupil size was sensitive to the internal state rather than simply reflecting visual stimulation. Similar to our list-level analyses (Figure 3B), the pupil size peaked at the second word presentation of the list and similar to our item-level analyses (Figure 3A), relative pupil size rose dramatically at the start of the distractor period. However, in contrast to the timing of our classifier predictions, which peak at 800–900 ms after word onset, pupil size peaks at 1–2 s after word onset. Further work is needed to establish a direct correspondence between these measures of brain activity and pupil size, and to establish the extent to which they reflect specific encoding processes or more general task engagement.

### Regional Contributions

Having found evidence that we can gauge encoding state from both single-item and list-level classifiers, we investigated how different patterns of activity contribute to successful encoding states. We found that spectral tilt across most regions was associated with better recall, similar to prior findings (Ezzyat et al., 2017). Interestingly though, the general brain-wide pattern of spectral tilt observed in item-level correlations was altered slightly in the list-level correlations, highlighting two key regions underlying memory encoding—the prefrontal cortex and hippocampus (Preston & Eichenbaum, 2013; Kuhl, Rissman, & Wagner, 2012). In the list-level correlations, while we observed spectral tilt patterns in association areas and especially in prefrontal cortex, we did not observe the same pattern in medial temporal areas, especially in hippocampus. This lack of spectral tilt in the hippocampus suggests that this region does not sustain its contribution to encoding state over multiple items.

Previous work has probed the idea of neural fatigue during encoding in the context of free recall, and in the process also illuminated regional differences in the dynamics of encoding state. Lohnas, Davachi, and Kahana (2020) tested the hypothesis that the hippocampus, which contributes to good encoding state through HFA (Sederberg et al., 2003; Long, Burke, & Kahana, 2014), can experience a depletion in neural resources that may then contribute to a poor encoding state. They compared hippocampal HFA during presentation of subsequently unrecalled items that followed a good encoding state (subsequently recalled items), to those that followed a similarly poor encoding state (subsequently unrecalled items), and found reduced HFA during items that followed good encoding states. In contrast to the hippocampus, which exhibited this evidence of neural fatigue, the DLPFC showed an opposite pattern of more persistent encoding state, where items following good encoding states had greater HFA than those following poor encoding states.

Building on these results, we hypothesized that these regional differences in neural fatigue would also relate to the temporal scale of hippocampal and DLPFC contributions to encoding state. Specifically, since the hippocampus fatigues at a fast rate of single items (as indexed by HFA), then hippocampal HFA should contribute to item-level encoding state more than list-level. Conversely, since DLPFC HFA exhibits a more persistent contribution to encoding, there HFA should contribute more to list-level than item-level state. Indeed, our results show exactly this pattern (Figure 6); univariate correlations between hippocampal HFA and recall performance are stronger at the item-level compared to the list-level, whereas correlations between DLPFC HFA and recall are stronger at the list-level. Although our results align with previous findings (Lohnas et al., 2020), elucidating their relation to neural fatigue would necessitate further analyses of the temporal dynamics of encoding-related activity.

Our findings and those from Lohnas et al. (2020) support the idea of complementary roles of the prefrontal cortex and hippocampus, in which new memories may shift from their reliance on hippocampal systems, to prefrontal systems, over time (Preston & Eichenbaum, 2013). They also support previous work suggesting a greater role for the prefrontal cortex in coding for coarse, compared to fine, temporal context (Jenkins & Ranganath, 2010). Besides contributing more to longer-term memory state, the prefrontal cortex is likely involved in modulation of other attention and memory systems during encoding (Reinhart, Woodman, & Posner, 2015). Thus, while the neural basis of the list-level encoding state overlaps significantly with the item-level state, they are also complementary, comprising different parts of a neural basis for memory encoding over longer time scales.

### Conclusions

The study of memory requires (often significant) delays between encoding of the memoranda and the memory assessment. The lack of explicit responses reflecting encoding success as it happens thus presents a challenge that researchers have sought to overcome through the use of implicit measures of brain activity. Our characterization of the temporal dynamics of brain states predicting subsequent memory performance complements previous work that strongly suggested that subsequent memory effects reflect endogenous processes related to encoding success rather than external factors that are correlated with recall performance (Urgolites et al., 2020; Weidemann & Kahana, 2021). We found that iEEG-based list-level classifiers of successful encoding perform on par with item-level classifiers, and likely reflect meaningful internal states of encoding and/or task engagement. Our work thus confirms the value of subsequent-memory analyses for the study of encoding processes, and opens up new possibilities for studying the associated dynamics.

---

**Figure S1.**
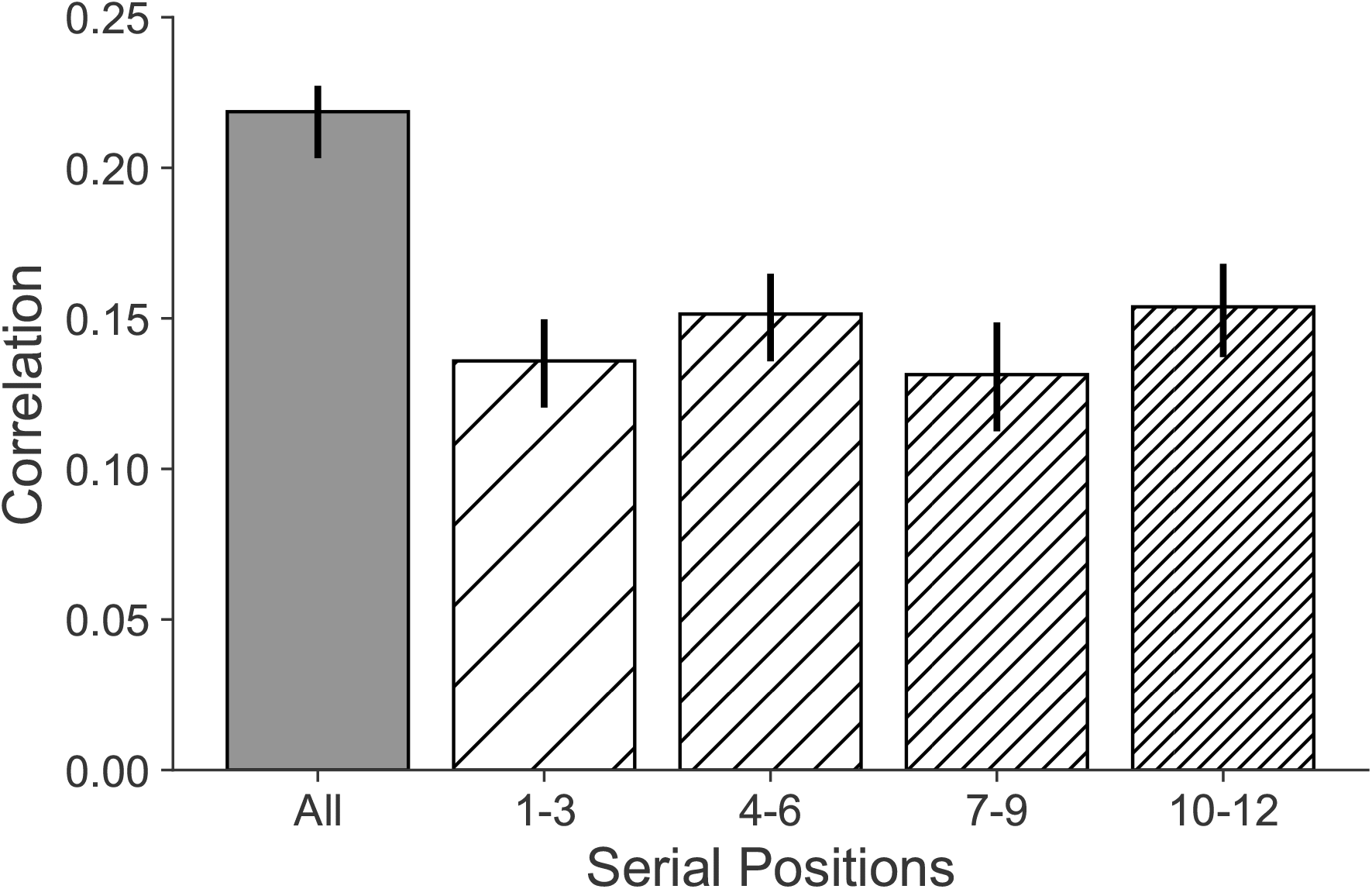
Serial Position Effect on Item-level Classification. We trained and tested item-level classifiers on subsets of the data, depending on serial position. For each serial position grouping, only items at those serial positions are included in the training set and testing set, and otherwise all other parameters are identical to the whole dataset. Classification performance is significantly greater for the whole set as compared with each of the four serial position subgroups (p’s < 0.001). Error bars indicate ± 1 SEM.

**Figure S2.**
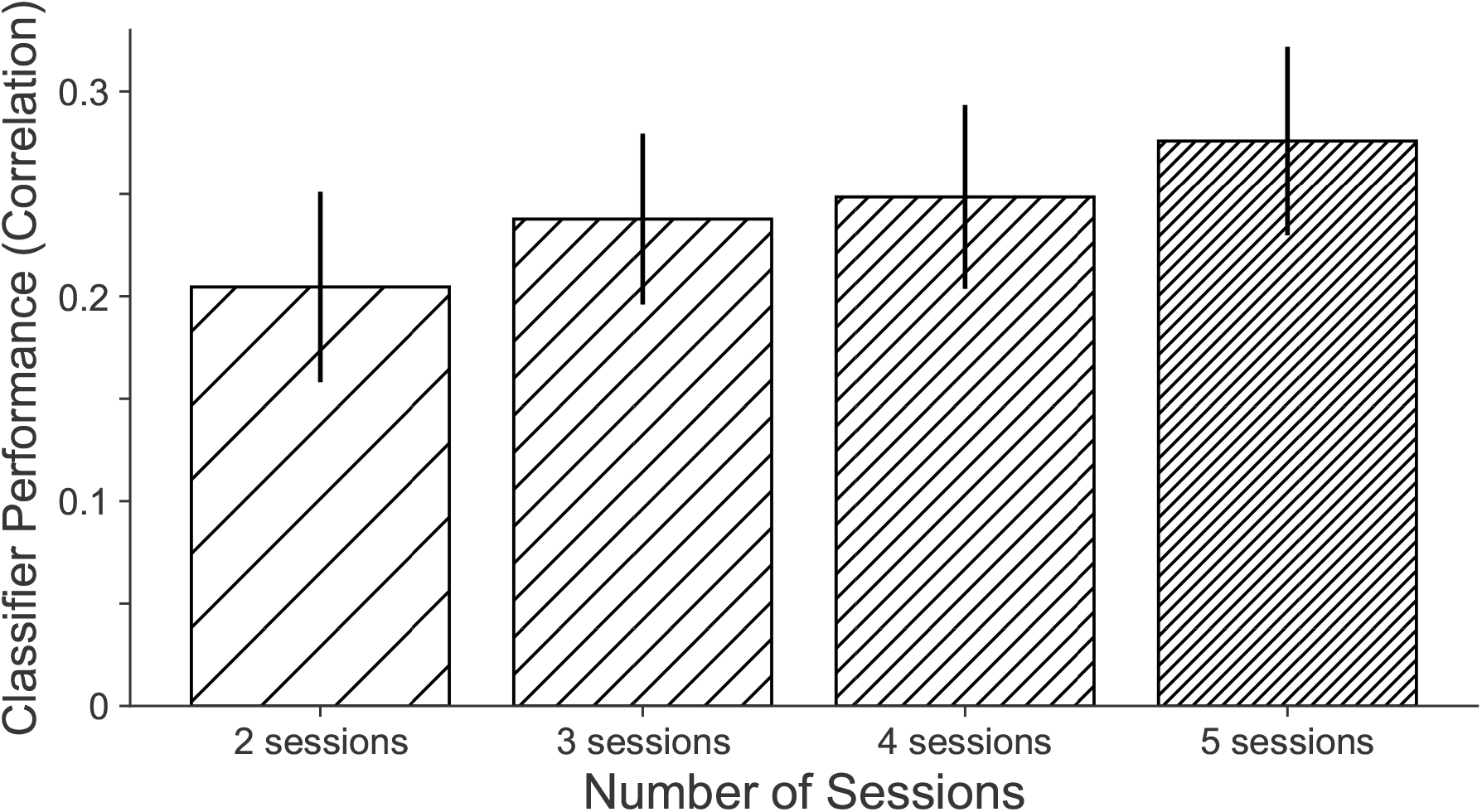
Classification performance increases with session number. In a subset of patients with at least five sessions of data (n = 22) we randomly selected two, three, four, or five sessions, 20 times. For each patient we evaluated classification performance by averaging the performance (correlation between predicted and observed list performances) of each iteration. Error bars represent ± 1 SEM.

**Figure S3.**
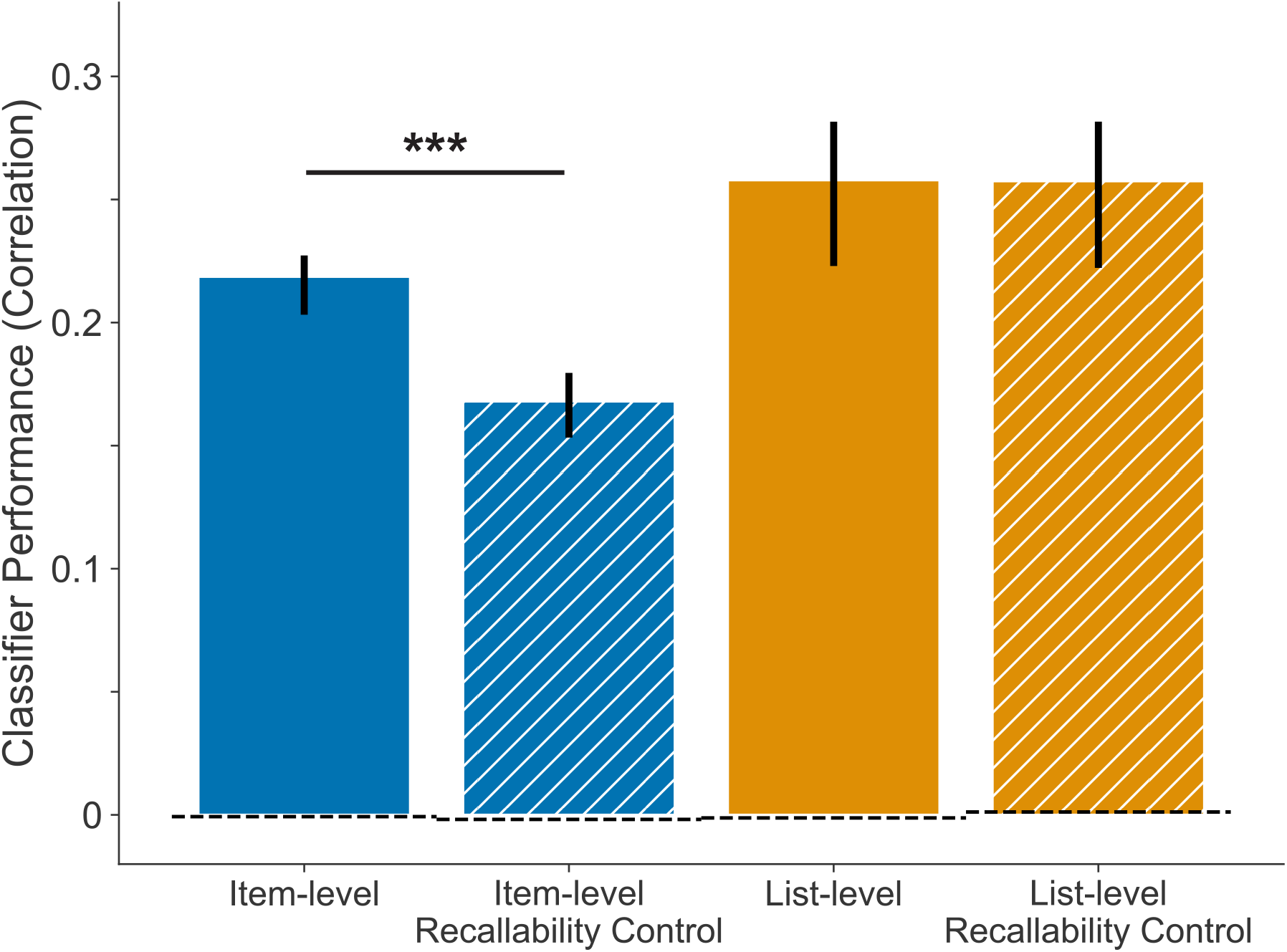
Classifier Performance with Recallability Correction. We quantified classifier performance as the correlation between predicted probability and observed recall (for item-level, blue) or predicted and observed list performance (for list-level, orange) after residualizing by overall recallability as measured based on all other subjects (hatched bars). We used permutation testing to derive the expected null correlation (dotted baselines), in which we randomly shuffled recall performances prior to classification for each subject (# permutations = 50). Error bars indicate ±1 SEM. *** denotes *p* < 0.001.

**Figure S4.**
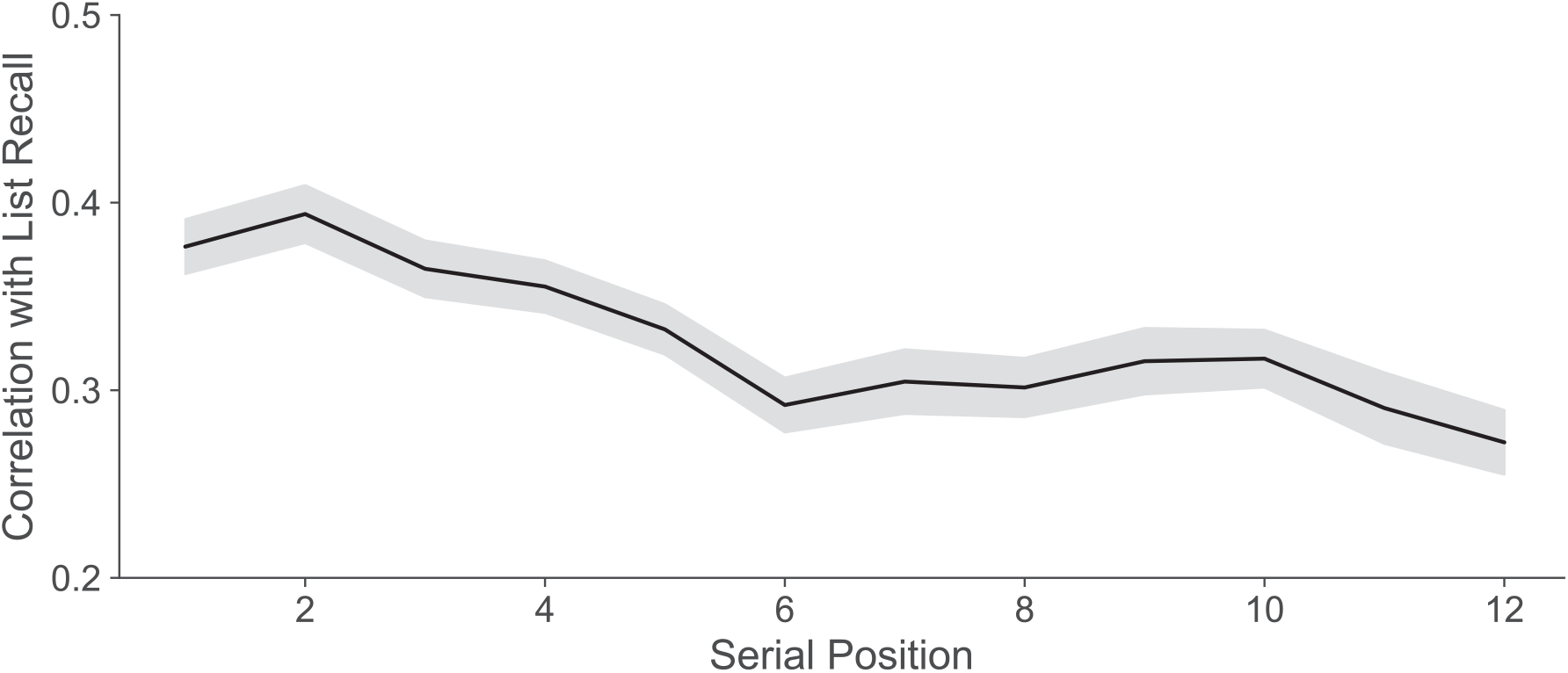
Correlation between Item Recall and List Performance by Serial Position. For each serial position, we plot the average correlation between list-level recall performance and recall of the items at that serial position. Despite *neural activity* of later serial positions being more predictive of list-level performance (Figure 4), *behaviorally-speaking*, recall of earlier serial positions is more predictive of list performance. Shaded error bars indicate ± 1 SEM.

**Figure S5.**
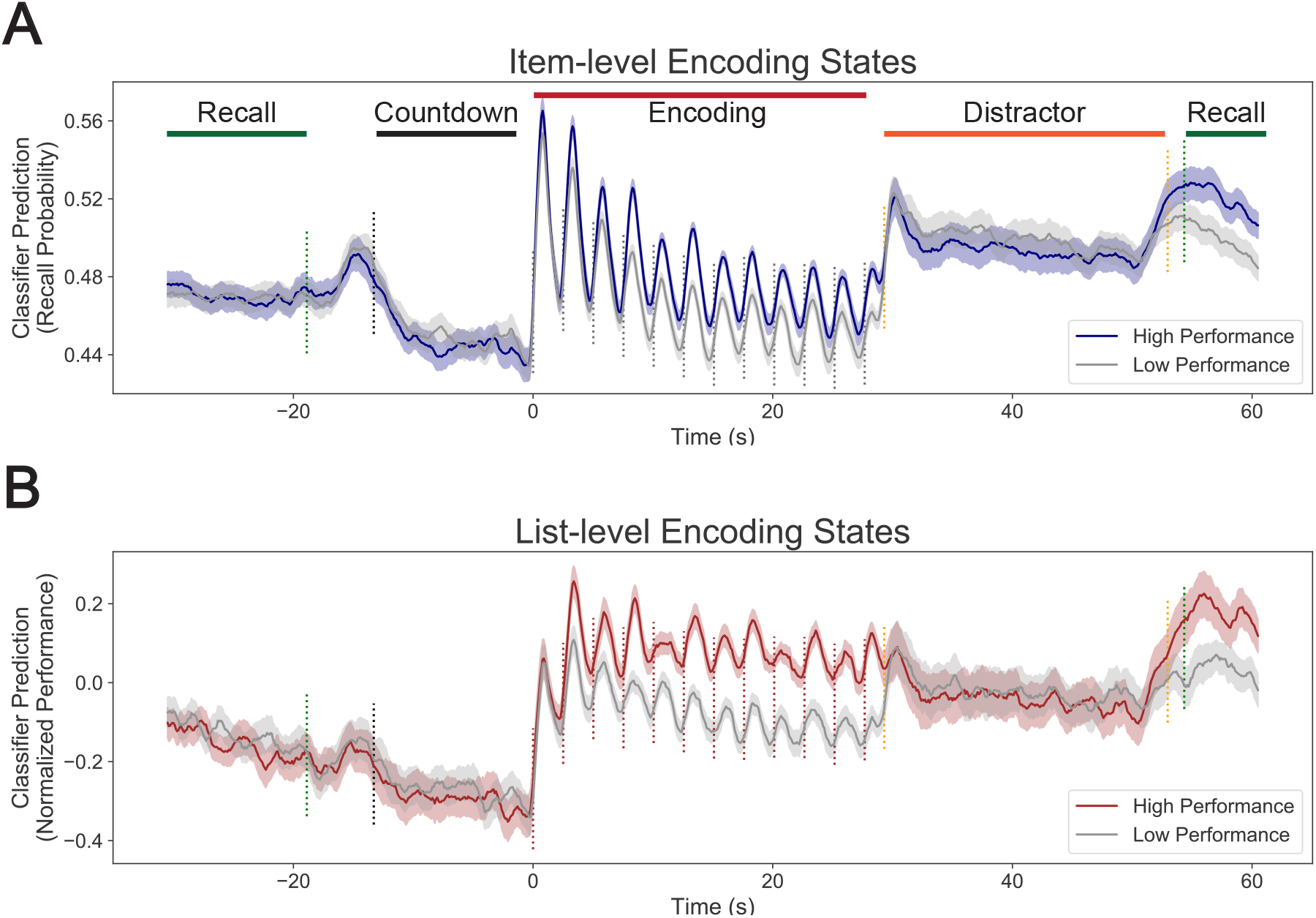
Classifier Predictions by Performance. (**A, B**) Time courses of item-level (A) and list-level (B) classifier predictions over the course of higher-performing lists (highest tertile performance, colored) and lower-performing lists (lowest tertile recall performance, gray), showing correspondence with task-relevant events and correspondence between classifier prediction and observed performance specifically during encoding and recall phases of the task, and not during distractor or countdown periods. We normalized list-level classifier prediction by subtracting the test list session’s mean performance. Dotted lines indicate average time of task events including start of countdown before list (black), word presentation times (brown), start/end of math distractor task (orange), and start/end of recall period (green). All shaded error regions indicate ± 1 SEM.

**Figure S6.**
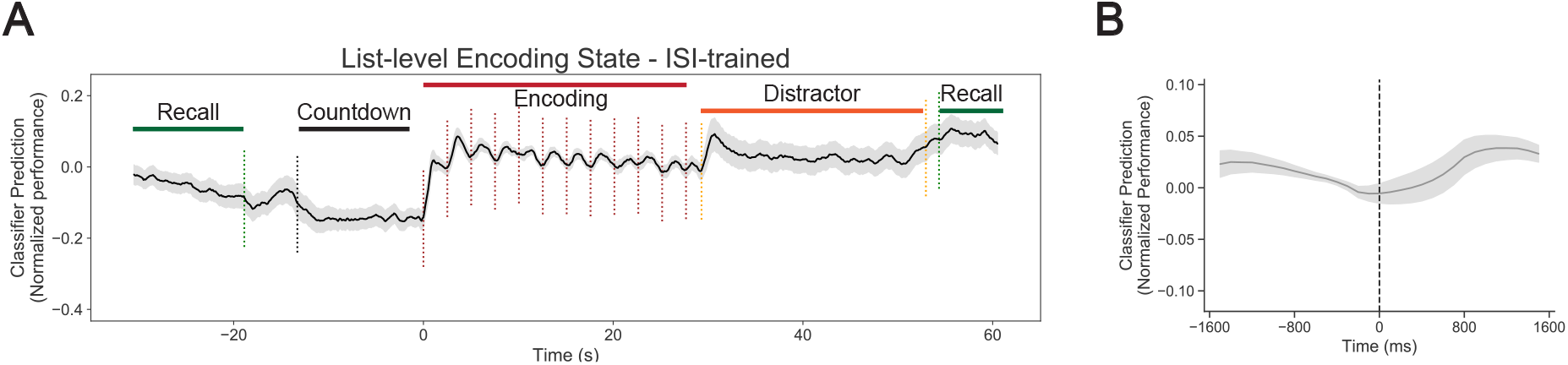
Classifier Predictions Based on ISI Training. (**A**) Time course of list-level classifier predictions over the course of encoding lists, showing correspondence with task-relevant events. Classifier is trained only on inter-stimulus intervals (700 ms preceding word presentations). We normalized list-level classifier prediction by subtracting the test list session’s mean performance. Dotted lines indicate average time of task events including start of countdown before list (black), word presentation times (brown), start/end of math distractor task (orange), and start/end of recall period (green). (**B**) Peri-stimulus time course of classifier predictions for list-level (A) classifier, time-locked to word presentation time, highlighting peak classifier predictions at 1200 ms. All shaded error regions indicate ± 1 SEM.

## References

Aka, A., Phan, T. D., & Kahana, M. J. (2021). Predicting Recall of Words and Lists. Journal of Experimental Psychology: Learning, Memory, and Cognition, 47(5), 765–784. doi: 10.1037/xlm0000964

Avants, B. B., Epstein, C. L., Grossman, M., & Gee, J. C. (2008). Symmetric diffeomorphic image registration with crosscorrelation: Evaluating automated labeling of elderly and neurodegenerative brain. Medical Image Analysis, 12(1), 26–41. doi: 10.1016/j.media.2007.06.004

Bainbridge, W. A., Hall, E. H., & Baker, C. I. (2019). Drawings of real-world scenes during free recall reveal detailed object and spatial information in memory. Nature Communications, 10(1), 5. doi: 10.1038/s41467-018-07830-6

Benjamini, Y., & Hochberg, Y. (1995). Controlling the False Discovery Rate: A Practical and Powerful Approach to Multiple Testing. Journal of the Royal Statistical Society: Series B (Methodological), 57(1), 289–300. doi: 10.1111/j.2517-6161.1995.tb02031.x

Burke, J. F., Sharan, A. D., Sperling, M. R., Ramayya, A. G., Evans, J. J., Healey, M. K., … Kahana, M. J. (2014). Theta and High-Frequency Activity Mark Spontaneous Recall of Episodic Memories. The Journal of Neuroscience, 34(34), 11355–11365. doi: 10.1523/JNEUROSCI.2654-13.2014

Corballis, M. C. (1969). Patterns of Rehearsal in Immediate Memory. British Journal of Psychology, 60(1), 41–49. doi: 10.1111/j.2044-8295.1969.tb01174.x

deBettencourt, M. T., Norman, K. A., & Turk-Browne, N. B. (2018). Forgetting from lapses of sustained attention. Psychonomic Bulletin and Review, 25(2), 605–611. doi: 10.3758/s13423-017-1309-5

Desikan, R. S., Ségonne, F., Fischl, B., Quinn, B. T., Dickerson, B. C., Blacker, D., … Killiany, R. J. (2006). An automated labeling system for subdividing the human cerebral cortex on MRI scans into gyral based regions of interest. NeuroImage, 31(3), 968–980. doi: 10.1016/j.neuroimage.2006.01.021

Donaldson, D. I., Petersen, S. E., Ollinger, J. M., & Buckner, R. L. (2001). Dissociating state and item components of recognition memory using fMRI. NeuroImage, 13(1), 129–142. doi: 10.1006/nimg.2000.0664

Ezzyat, Y., Kragel, J. E., Burke, J. F., Levy, D. F., Lyalenko, A., Wanda, P., … Kahana, M. J. (2017). Direct Brain Stimulation Modulates Encoding States and Memory Performance in Humans. Current Biology, 27(9), 1251–1258. doi: 10.1016/j.cub.2017.03.028

Gramfort, A., Luessi, M., Larson, E., Engemann, D. A., Strohmeier, D., Brodbeck, C., … Hämäläinen, M. (2013). MEG and EEG data analysis with MNE-Python. Frontiers in Neuroscience, 7, 267. doi: 10.3389/fnins.2013.00267

Griffiths, B., Mazaheri, A., Debener, S., & Hanslmayr, S. (2016). Brain oscillations track the formation of episodic memories in the real world. NeuroImage, 143, 256–266. doi: 10.1016/j.neuroimage.2016.09.021

Halpern, D. J., Tubridy, S., Davachi, L., & Gureckis, T. M. (2021). Identifying Causal Subsequent Memory Effects. bioRxiv, 1–31. Retrieved from https://www.biorxiv.org/content/early/2021/11/10/2021.11.08.467782 doi: 10.1101/2021.11.08.467782

Hill, P. F., King, D. R., Lega, B. C., & Rugg, M. D. (2020). Comparison of fMRI correlates of successful episodic memory encoding in temporal lobe epilepsy patients and healthy controls. NeuroImage, 207, 116397. doi: 10.1016/j.neuroimage.2019.116397

Jenkins, L. J., & Ranganath, C. (2010). Prefrontal and Medial Temporal Lobe Activity at Encoding Predicts Temporal Context Memory. The Journal of Neuroscience, 30(46), 15558–15565. doi: 10.1523/JNEUROSCI.1337-10.2010

Kahana, M. J., Aggarwal, E. V., & Phan, T. D. (2018). The Variability Puzzle in Human Memory. Journal of Experimental Psychology: Learning, Memory, and Cognition, 44(12), 1857–1863. doi: 10.1037/xlm0000553

Kim, H. (2011). Neural activity that predicts subsequent memory and forgetting: A meta-analysis of 74 fMRI studies. NeuroImage, 54(3), 2446–2461. doi: 10.1016/j.neuroimage.2010.09.045

Kucewicz, M. T., Dolezal, J., Kremen, V., Berry, B. M., Miller, L. R., Magee, A. L., … Worrell, G. A. (2018). Pupil size reflects successful encoding and recall of memory in humans. Scientific Reports, 8(1), 4949. doi: 10.1038/s41598-018-23197-6

Kuhl, B. A., Rissman, J., & Wagner, A. D. (2012). Multi-voxel patterns of visual category representation during episodic encoding are predictive of subsequent memory. Neuropsychologia, 50(4), 458–469. doi: 10.1016/j.neuropsychologia.2011.09.002

Lohnas, L. J., Davachi, L., & Kahana, M. J. (2020). Neural fatigue influences memory encoding in the human hippocampus. Neuropsychologia, 143, 107471. doi: 10.1016/j.neuropsychologia.2020.107471

Long, N. M., Burke, J. F., & Kahana, M. J. (2014). Subsequent memory effect in intracranial and scalp EEG. NeuroImage, 84, 488–494. doi: 10.1016/j.neuroimage.2013.08.052

Merkow, M. B., Burke, J. F., Stein, J. M., & Kahana, M. J. (2014). Prestimulus Theta in the Human Hippocampus Predicts Subsequent Recognition But Not Recall. Hippocampus, 24(12), 1562–1569. doi: 10.1002/hipo.22335

Murdock, B. B. (1962). The serial position effect of free recall. Journal of Experimental Psychology, 64(5), 482–488. doi: 10.1037/h0045106

Otten, L. J., Henson, R. N., & Rugg, M. D. (2002). State-related and item-related neural correlates of successful memory encoding. Nature Neuroscience, 5(12), 1339–1344. doi: 10.1038/nn967

Paller, K. A., & Wagner, A. D. (2002). Observing the transformation of experience into memory. Trends in Cognitive Sciences, 6(2), 93–102. doi: 10.1016/S1364-6613(00)01845-3

Park, H., & Rugg, M. D. (2010). Prestimulus hippocampal activity predicts later recollection. Hippocampus, 20(1), 24–28. doi: 10.1002/hipo.20663

Preston, A. R., & Eichenbaum, H. (2013). Interplay of Hippocampus and Prefrontal Cortex in Memory. Current Biology, 23(17), R764–R773. doi: 10.1016/j.cub.2013.05.041

Reinhart, R. M., Woodman, G. F., & Posner, M. I. (2015). Enhancing long-term memory with stimulation tunes visual attention in one trial. Proceedings of the National Academy of Sciences, USA, 112(2), 625–630. doi: 10.1073/pnas.1417259112

Sederberg, P. B., Kahana, M. J., Howard, M. W., Donner, E. J., & Madsen, J. R. (2003). Theta and Gamma Oscillations during Encoding Predict Subsequent Recall. The Journal of Neuroscience, 23(34), 10809–10814. doi: 10.1523/JNEUROSCI.23-34-10809.2003

Serruya, M. D., Sederberg, P. B., & Kahana, M. J. (2014). Power Shifts Track Serial Position and Modulate Encoding in Human Episodic Memory. Cerebral Cortex, 24(2), 403–413. doi: 10.1093/cercor/bhs318

Stevens, S., Valderas, J. M., Doran, T., Perera, R., & Kontopantelis, E. (2016). Analysing indicators of performance, satisfaction, or safety using empirical logit transformation. BMJ, 352, i1114. doi: 10.1136/bmj.i1114

Tulving, E., & Rosenbaum, R. S. (2006). What do explanations of the distinctiveness effect need to explain? In R. R. Hunt & J. B. Worthen (Eds.), Distinctiveness and memory (pp. 406–423). New York, NY: Oxford University Press. doi: 10.1093/acprof:oso/9780195169669.003.0018

Urgolites, Z. J., Wixted, J. T., Goldinger, S. D., Papesh, M. H., & Treiman, D. M. (2020). Spiking activity in the human hippocampus prior to encoding predicts subsequent memory. Proceedings of the National Academy of Sciences, USA, 117(24), 13767–13770. doi: 10.1073/pnas.2001338117

Wagner, A. D., Schacter, D. L., Rotte, M., Koutstaal, W., Maril, A., Dale, A. M., … Buckner, R. L. (1998). Building Memories: Remembering and Forgetting of Verbal Experiences as Predicted by Brain Activity. Science, 281(5380), 1188–1191. doi: 10.1126/science.281.5380.1188

Weidemann, C. T., Kadel, A., Davis, K. A., Kragel, J. E., Wanda, P. A., Worrell, G. A., … Rizzuto, D. S. (2019). Neural Activity Reveals Interactions Between Episodic and Semantic Memory Systems During Retrieval. Journal of Experimental Psychology: General, 148(1), 1–12. doi: 10.1037/xge0000480

Weidemann, C. T., & Kahana, M. J. (2021). Neural Measures of Subsequent Memory Reflect Endogenous Variability in Cognitive Function. Journal of Experimental Psychology: Learning, Memory, and Cognition, 47(4), 641–651. doi: 10.1037/xlm0000966

